# Resolving when (and where) the Thylacine went extinct

**DOI:** 10.1101/2021.01.18.427214

**Authors:** Barry W. Brook, Stephen R. Sleightholme, Cameron R. Campbell, Ivan Jarić, Jessie C. Buettel

## Abstract

Like the Dodo and Passenger Pigeon before it, the predatory marsupial Thylacine (*Thylacinus cynocephalus*), or ‘Tasmanian tiger’, has become an iconic symbol of human-caused extinction. The last captive animal died in 1936, but even today reports of the Thylacine’s possible ongoing survival in remote regions of Tasmania are newsworthy and capture the public’s imagination. Extirpated from mainland Australia in the mid-Holocene, the large island of Tasmania became the species’ final stronghold. Following European settlement in the 1800s, the Thylacine was heavily persecuted and pushed to the margins of its range, although many sightings were reported thereafter—even well beyond the 1930s. To gain a new depth of insight into the extinction of the Thylacine, we assembled an exhaustive database of 1,237 observational records from Tasmania (from 1910 onwards), quantified their uncertainty, and charted the patterns these revealed. We also developed a new method to visualize the species’ 20^th^-century spatio-temporal dynamics, to map potential post-bounty refugia and pinpoint the most-likely location of the final persisting subpopulation. A direct reading of the high-quality records (confirmed kills and captures, in combination with sightings by past Thylacine hunters and trappers, wildlife professionals and experienced bushmen) implies a most-likely extinction date within four decades following the last capture (i.e., 1940s to 1970s). However, uncertainty modelling of the entire sighting record, where each observation is assigned a probability and the whole dataset is then subject to a sensitivity analysis, suggests that extinction might have been as recent as the late 1980s to early 2000s, with a small chance of persistence in the remote south-western wilderness areas. Beyond the intrinsically fascinating problem of reconstructing the final fate of the Thylacine, the new spatio-temporal mapping of extirpation developed herein would also be useful for conservation prioritization and search efforts for other rare taxa of uncertain status.

## Introduction

Prior to European settlement in the early 1800s (Jeffreys 1820), the large island of Tasmania supported a small but stable endemic population of a cursorial predator called the Thylacine (*Thylacinus cynocephalus*)—also known as the ‘marsupial wolf’ or ‘Tasmanian tiger’ (Guiler 1985)—that had become extirpated on mainland Australia during the late Holocene, after surviving the earlier wave of late Pleistocene megafaunal extinctions (Prowse et al. 2014). Due to deliberate persecution (encouraged by government and private bounties, paid out from 1888 to 1909), incidental snaring and trapping by fur traders, occasional capture for the zoo trade, habitat modification and possibly disease (Paddle 2012), the Thylacine had declined to extreme rarity by the early 20th century. The final confirmed wild captures and kills of the species occurred in the 1930s (Sleightholme & Campbell 2016). Recent modelling using annual sighting data has concluded that extinction occurred soon thereafter (Carlson et al. 2018)

The last captive Thylacine died in the Hobart Zoo on 7th September 1936, a date now commemorated annually as ‘Threatened Species Day’ in Australia. Fifty years later, in 1986, the species was formally designated as Extinct by the International Union for the Conservation of Nature. However, with many unconfirmed sightings reported in the decades after the 1930s (Rounsevell & Smith 1982), speculation has run rife that the Thylacine might have persisted far longer than formally accepted in the wilderness of Tasmania (Drollette 1996). Indeed, as one of the most famous of recently ‘extinct’ species, and an archetype of convergent evolution (with placental canids) (Owen 2003), the details surrounding the final fate of the Thylacine in its last island stronghold is fascinating to both the public and conservation science (Bulte et al. 2003).

Complicating matters, apparently reliable sightings came from former trappers and bushmen through to the 1960s (after which most had long retired or died). Dedicated expeditions continued thereafter, including an intense localized search by authorities in 1982 following a highly rated sighting by a National Parks officer (Guiler & Godard 1998). The regularity and frequency of apparently plausible but unverified sightings reported over the last 85 years has not only raised the Thylacine to iconic status in the global public’s eye, but also made it paradoxically challenging to reconstruct the timeline of its fate scientifically. Past efforts to prove the ongoing persistence of the Thylacine involved deliberate and sustained (albeit geographically restricted) field searches (Griffith 1972; Smith 1981), with them sometimes being financially motivated (e.g., media mogul Ted Turner offered a prize of $100,000 in 1983, and *The Bulletin* magazine of $1.25 million in 2005). More recently, mathematical models have been used to estimate the extinction time, but the scope of their inferences have—to date—been hampered by lack of data, being based on a simplified sighting record (one observation per year) or using generalizations from life-history correlates and analogies with other taxa (Lee et al. 2017; Brook et al. 2018). Although statistical approaches for extinction inference using a time series of observational records have been developed to incorporate observational quality explicitly (Boakes et al. 2015; Kodikara et al. 2018), these could only be applied to the Thylacine’s case in a highly constrained way (i.e., by designating each year as either a certain, uncertain or no record: Carlson et al. 2018), because the sighting data and the circumstances surrounding each reported incident had never been systematically collated or quality rated. Given the inherent complexities and data constraints, what can be said, scientifically, about the extinction date of the Thylacine? To tackle this intriguing problem, we compiled a comprehensive, quality-rated database of post-bounty Thylacine sighting records from Tasmania and devised a spatio-temporal method to visualize its extinction dynamics. The sighting database was amassed by searching and cataloguing records from official government archives, published reports, museum collections, newspaper articles, microfilm, contemporary correspondence, private collections or other miscellaneous citations and testimony. Each observation was dated, geotagged, quality-rated, categorized by type, and linked to image files of the original source material. Only post-1910 records were considered, being the period following the government bounty after which the species was considered rare (Guiler 1985). Records were classified as physical specimens, expert sightings, other observations, signs (e.g., tracks), and were all rigorously quality rated.

Using this curated sighting database, our goal was to evaluate alternative scenarios for the timing of extinction of the Thylacine and mapped the pattern of its preceding regional extirpations. To do this, we applied a recently developed frequentist method for inferring the probability of persistence at any date beyond that of the final record under circumstances where many observations are uncertain and not equally reliable (Brook et al. 2019). This combines a probabilistic re-sampling of sighting records with a statistical extinction-date estimator (EDE), focusing on two models: i) the optimal linear estimator (OLE), used for famous examples such as the Dodo (*Raphus cucullatus)* (Roberts & Solow 2003), and ii) the variable-sighting-rate method (McInerny et al. 2006). Both EDE models are robust to cases where sighting frequency declines over time (Jarić & Roberts 2014). In this context, we considered a wide range of viewpoints and assumptions, from the conservative (i.e., rejection of all unverified records), to the liberal (i.e., use of all reported sightings, weighted by sighting quality). Our aim was not to determine precisely when (and where) the last Thylacine died, but to offer plausible scenarios.

## Methods

The approach involved the following steps: (i) collation and georeferencing of Thylacine sighting records from Tasmania into a database of unique observations; (ii) error-checking, attribute scoring, and quality-rating; (iii) a scenario analysis, using EDEs coupled with selective rejection or weighted selection of sighting records, to infer the island-wide and regional distribution of times to extinction for the Thylacine; (iv) a sensitivity analysis on the impact on extinction time of bioregional disaggregation and different record-inclusion probabilities; and (v) dynamic distribution mapping, to illustrate the spatial patterns of extirpation.

### Database compilation and validation

We compiled and error-checked a comprehensive repository of documented Thylacine sighting records from Tasmania, covering the period from the year after the last Government bounty was paid (1910) to the present. We refer to this as the Tasmanian Thylacine Sighting Records Database (TTSRD). By examining sources exhaustively, spanning official archives, published reports, past (partial) compilations of sighting records (Smith 1981; Rounsevell & Smith 1982; Guiler 1985), museum collections, newspaper and media articles, microfilm, contemporary correspondence, private collections and other miscellaneous citations and testimony, we were able to amass 1,237 unique observations from this period, as well as resolve previous anomalies and duplications.

The TTSRD is presented in a flat-file format (.xlsx or .csv), with one observation per row, and column data for a unique ID, sighting location (with notes), date (year/month [or season]), geo-reference (latitude/longitude and a precision class), sighting type (kill, capture, expert or non-expert visual observation, secondary evidence) and quality-rating (a score between 1 [lowest] to 5 [highest], based on a subjective composite of information regarding the observer’s credentials and experience, number of observers, context, and veracity of the description supplied), sighting meta-data (road or bush, driving or walking, day or night, near or far distance, number of observers, and number of Thylacines recorded), observer remarks/notes, and a reference and link to an image of its source material(s). Confidentiality requirements necessitated the redaction of names and addresses for 89 records, but efforts were made to prevent this censure affecting the essential content of the reports. The TTSRD was checked rigorously for repeated or erroneous entries (e.g., there were examples when the year or location did not exactly match, or when observer’s names were misspelled, but other corroborating or correlative evidence pointed to a duplication). Any errors we found were archived and removed, with the record’s unique ID not reused.

### Extinction date estimation

Given the difficulties inherent in observing rare or critically endangered taxa, statistical methods (extinction dynamics estimators: EDE) have been developed to *infer* probable extinction dates (or, by inversion, probabilities of persistence) from a time series of sighting records. Recent approaches to EDE have sought to incorporate a mixture of certain and uncertain sighting records (Boakes et al. 2015). Both Bayesian (Solow & Beet 2014; Kodikara et al. 2018) and frequentist (Jarić & Roberts 2014; Brook et al. 2019) methods have been developed, each with advantages, limitations and differing philosophical framings of the problem.

In this analysis, because of the character of the data embodied within the TTSRD (see Results), we chose to use a recently developed frequentist method for the inclusion and relative weighting of observations (Jarić & Roberts 2014; Brook et al. 2019), implemented in Program R (v4.1), because this approach can incorporate explicit sighting probabilities on uncertain data, multiple sightings within a year, and mixed records of variable type and quality. The approach makes use of two established frequentist EDE (Roberts & Solow 2003; McInerny et al. 2006) for statistical inference, but probabilistically re-samples the observations of the full sighting record to generate a frequency distribution of extinction times, as demonstrated and validated using various example species records in Brook et al. (2019). Note that in all frequentist EDE models, the null hypothesis underlying the statistical inference is that of persistence, with each sighting assigned a probability of being correct, given an assumption that the species persisted up to at least the point in time when the observation was made. A detailed evaluation of the assumptions underpinning this approach, and the alternatives, are given in the Supporting Information, Appendix S2.

### Scenarios and sensitivity analysis for range-wide extinction

Our analysis is predicated, in large part, on assumptions about which records are true and which are false (whether from misidentification, illusion or deception). In terms of possible scenarios, the most conservative assumption is to accept only those records based on kills or captures, in which a body, a photograph of a body, or a live animal was produced, i.e., physical specimens. We first used standard EDEs to model these records, using either island-wide or regional collections. We then experimented with the remaining dataset of uncertain observations, to assess the impact of applying different inclusion/exclusion criteria, in concert with the probabilistic weightings, on the extinction-year estimate (the physical records are included in all other scenarios because they are certain and should never be rejected). For example, one possibility is to also include unconfirmed kills and captures (where a kill was reported but the body was left behind, or an animal was trapped but then released or it accidentally escaped during handling). Another is to set a plausibility threshold on uncertain sightings, accepting only those that meet rigorous quality standards, and rejecting all others, or to only consider sightings that were made by two or more witnesses. Similarly, a threshold could be applied to a date, by accepting records prior to a given year and rejecting those reported afterwards (in reality, after the true extinction year, all sightings are axiomatically false; however, this year is of course not known, *a priori,* being the variable under question). Finally, a mixed-certainty EDE could be applied, using some (e.g., only ‘exper-trated’ sightings), or all records, with each sighting assigned a probability of being true, and scenarios constructed based on different relative weightings. We tried examples of all these approaches herein, while acknowledging that there is essentially no limit to the alternative interactions and dependencies among assumptions that might be imagined.

A sensitivity analysis was also done on inferences made by the EDE-model-under-uncertainty using all records (reported in the Results), with additional variants, such as alternative record-weighting schemes, and alternative EDE parameterizations, reported in the Supporting Information, Appendix S4. Therein, Table S3 gives the relative probability (P) weightings of records by type (physical specimen, expert observation, expert indicator of presence, and other observations or indicators), for low (L), default (D) and high (H) weights, and Table S4 reports the probability multipliers on the weightings given in Table S3, which depend on the quality rating. Table S5 shows the results of scenarios for the date of extinction of the Thylacine in Tasmania, based on all sightings (rather than physical specimens and expert sightings only). Extinction-date inferences are also given for individual bioregions, based on the combination of physical specimens and expert-sighting records (Table S6), and for all available records (Table S7).

### Pattern of regional extirpation

Both the last capture and the last confirmed kill of a Thylacine in the wild was situated in a semi-agricultural region of north-west Tasmania (Sleightholme et al. 2020). However, many reports continued to come thereafter from more remote central and south-west regions of the island, a vast stretch of wilderness that was sparsely settled and relatively rarely traversed or trapped. To differentiate regional extirpations (as a nearly inevitable prelude to global extinction), we used the Interim Biogeographic Regionalization of Australia (IBRA7) framework (Department of the Environment 2012) as a basis for disaggregating the TTSRD records spatially into ecologically distinct Tasmanian bioregions. These spatial units are defined based on common climate, geology, landform, native vegetation, and species information.

This biogeographic division of the TTSRD has the advantage of permitting a semi-spatial breakdown of the sighting records into an ecologically meaningful regionalization—suitable for discrimination of discrete-spatial patterns of extirpation—whilst still yielding enough records for a statistically robust EDE-based inference. A limitation of this approach of biogeographic analysis is that it assumes implicitly that the sub-populations of Thylacine across regions follow independent fates (i.e., they are not interconnected by dispersal). In practice, although bioregional delineations like IBRA7 are intended to be ecologically consistent, their boundaries typically do not constitute landscape-scale barriers.

For a broader spatial perspective on extirpation, we also tried aggregating the smaller bioregions into four larger, spatially coherent clusters (*South-East* = Tasmanian Northern Midlands, Tasmanian South East; *North-East* = Ben Lomond, Furneaux; *North-West* = Tasmanian Northern Slopes, King; *West / World Heritage Area* = Tasmanian Central Highlands, Tasmanian West, Tasmanian Southern Ranges) and analyzed the Thylacine record collections separately for each.

### Spatially continuous mapping of extirpation

To create a visualization of the extirpation pattern and relax the ‘hard-boundary’ assumption inherent in the approach based on bioregional divisions, we developed a spatially continuous mapping method, implemented algorithmically in R. The input is a raster map of grid cells representing the landscape occupied by the species. In our case, we gridded the main island of Tasmania into 69,562 × 0.01° longitude-latitude cells (~ 0.92 km^2^ at the Tasmanian latitudinal range of −40.65 to −43.64° S). For each grid cell, a subset of sighting records within a pre-defined radius are selected (we chose a threshold of 75 km for the Thylacine in Tasmania). Sighting records within that subset are then weighted for inclusion in the probabilistic EDE based on a distance-decay function. We used a truncated exponential model, *w* = min(1, *ab^-d^*), where *a* = 2.15, *b* = 1.074 and *d* is the distance (in km) from the target cell (other mathematical models could be used, if deemed more appropriate to a situation). These values were used because the model parameterized in this way led to a weighting (*w*) of 1 for all records within 10 km distance, declining to approximately 0.5 by 20 km, 0.25 by 30 km, 0.05 by 50 km, and 0.01 by 75 km, which is appropriate for what we know of the Thylacine’s probable home range and dispersal patterns (Guiler 1985). For use in the EDE, the *w* value of each record is multiplied by its sighting probability (see above, ‘Extinction date estimation’ section). Because of the large number of grid cells, we used the variable sighting rate EDE model (McInerny et al. 2006), because an analytical form of the under-record-selection-uncertainty framework has been derived (see Jarić & Roberts 2014); this method was used because it is more computationally efficient than the generalized (but slower) re-sampling approach. When iterated over all spatial-grid cells, this algorithm produces a final map of inferred extirpation times. For enhanced execution speed, the R code makes use of multi-core parallel processing.

The obvious advantage of a spatially continuous approach is that it generates a smoothed geographic surface and contours of extirpation times that can be mapped, based on the inferred year of extinction or upper confidence bound, as derived from an EDE. Importantly, because the method makes use of all records within a defined radius (with their selection probability down-weighted monotonically by distance), it does not impose hard boundaries like in the discrete-bioregional approach and captures spatial-pattern information from both within and across the bioregions. Moreover, regions of high uncertainty (wide confidence bounds) are readily distinguished (and visualized) from those underpinned by more data and/or better-constrained inferences on extirpation. The main limitation is that the mathematic form (and parameter values) of the distance-decay function is ultimately subjective, yet these choices govern the degree of spatial smoothing apparent in the resultant probability map. Although the spatially continuous mapping algorithm was developed specifically for the Thylacine, the approach (and R code) is designed to be general and could be readily applied to other species with geo-referenced sighting records.

## Results

### Database metrics

The final database comprised 1,237 entries (99 physical records, 429 expert sightings), with observations from all years except 1921, 2008 and 2013. Many records from 1910 to 1936 (the year the last captive specimen died—a male captured in 1931 (Sleightholme et al. 2020), see photograph in Fig. 1) were of confirmed kills or live captures, although 56.6 % (128) of the 226 entries dating from this period were unverified sightings. The last fully documented wild animal (with photographs) was shot in 1930, but there is little reason to doubt the legitimacy of two bodies noted from 1933, nor two other capture-and-releases from 1935 and 1937. Thereafter, over the course of eight decades, a further 26 deaths and 16 captures were reported (but not verified), along with 271 sightings by ‘experts’ (e.g., former trappers, bushmen, scientists or officials). The remaining 698 observations from Tasmania were made by the general public.

**Fig. 1.**
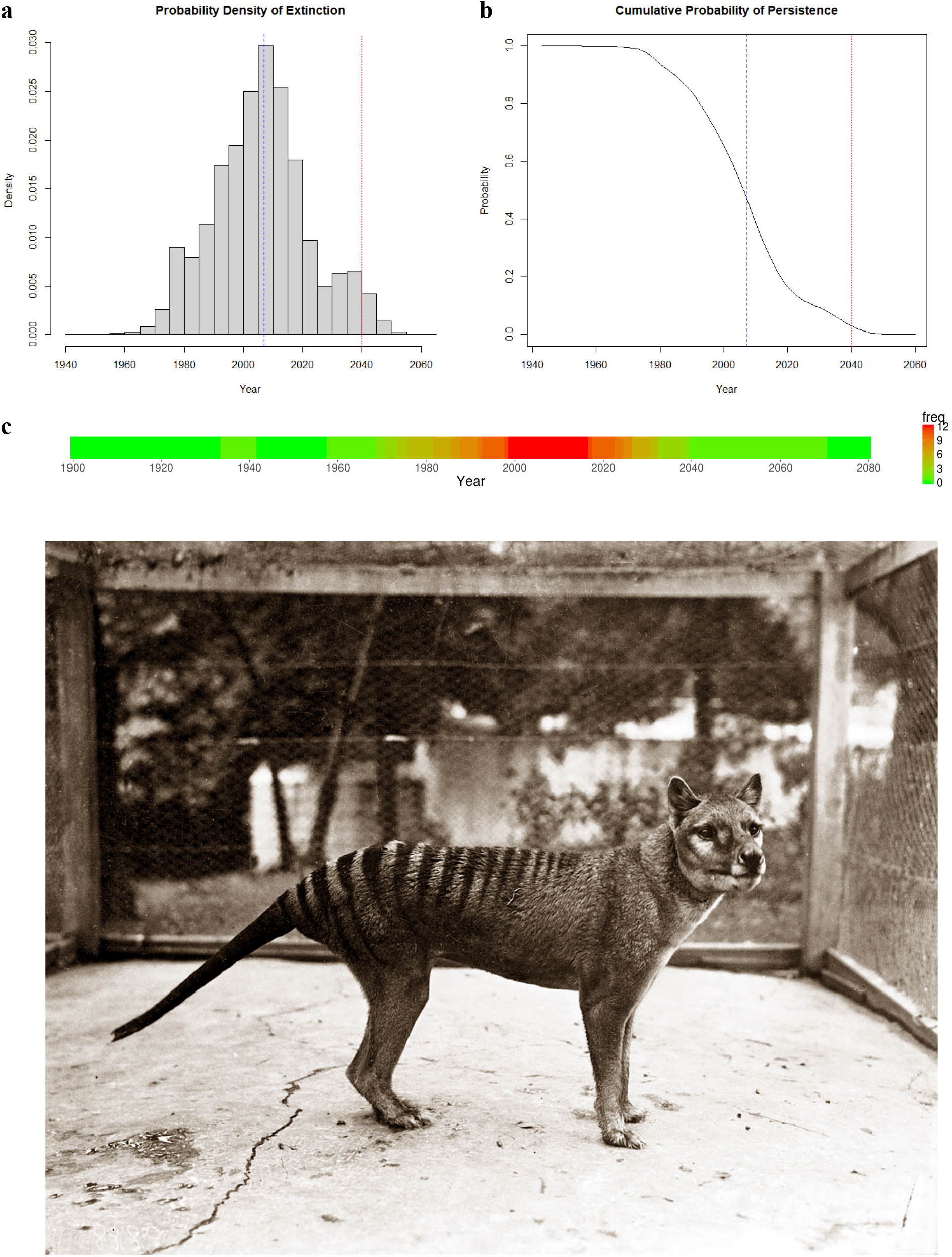
Simulated extinction dates for the Thylacine in Tasmania, using all 1,237 quality-rated sighting records. **a**. Probability-density distribution of the inferred extinction date from the optimal linear estimator, based on probabilistic re-sampling of all 1,237 specimens and observational records from 1910–2019, with the low scenario for probability weightings on the uncertain records. **b**. Cumulative probability of persistence at a given calendar year, as derived from the distribution shown in **a**. In each panel, the blue and red vertical lines show the mean time of extinction and upper 95% confidence bound, respectively. **c**. Sensitivity heatmap, a merger of upper/lower -bound weights assigned to the sighting-type probabilities (default/conservative): physical records = 1/1, expert observations = 0.25/0.05, expert indications (e.g., footprints, scats) = 0.1/0.01, other observations = 0.05/0.005, other indications = 0.01/0.001. Photograph is of the last captive Thylacine, taken on 19th December 1933 at the Hobart Zoo by zoologist David Fleay (image courtesy David Fleay trustees).

There were notable spikes in reporting rates in 1937 and 1970, the former following legal protection and the latter arising from media attention linked to a well-publicized expedition. There are also many examples of discrete spatio-temporal sighting clusters with closely matching visual descriptions, the interrelationships of which would not have been apparent at the time the reports were submitted to authorities. Overall, the annual number of reports in the six decades spanning 1940–1999 were relatively constant 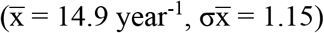, but fell substantially 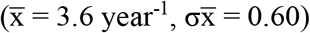 from 2000–present. A breakdown of observations by type and quality, and time-series plots, are reported in the Supporting Information (Appendix S4).

### Range-wide extinction scenarios and sensitivity analysis

Restricting the EDE inference solely to the physical specimens results in an uncontroversial conclusion: extinction by 1941, with a regional east-to-west pattern of loss (Table 1). The physical records ceased earliest in the midlands and south-east of the island, which coincides with where human settlements and farming were most widespread in the early 20^th^ century. If the uncertain kills and captures are also considered, the lower-bound for extinction is pushed out to the mid-1950s, with the median estimate spanning the 1970s to 1980s (depending on which EDE is used). If only the highest-quality uncertain sightings are included alongside the physical records, then the lower-bound is the late-1950s to early 1960s, with the median estimates spanning the late 1980s to 1990s; similar results come from only considering sightings that were reported by multiple witnesses (Table 2a). Arbitrarily fixing a cut-off date does not seem helpful, because the sudden truncation of records causes both EDE models to project extinction within a year of the applied threshold, leading to a somewhat circular result.

**Table 1.**
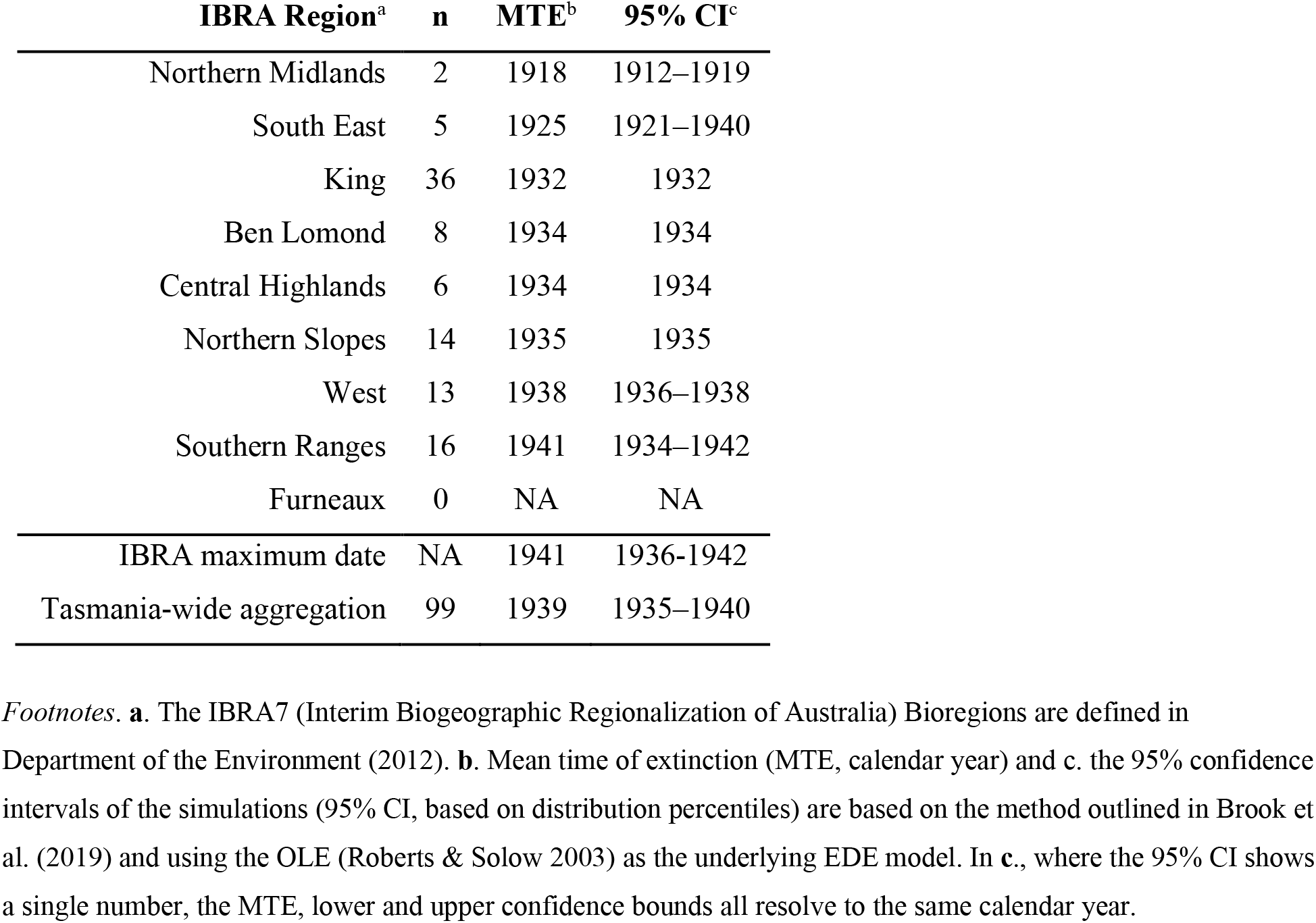
Regional and island-wide extinction date inference for the Thylacine in Tasmania, based solely on physical records (i.e., excluding unconfirmed sightings).

**Table 2.** Scenarios for the date of extinction of the Thylacine in Tasmania.

|  |  |  | Variable Sighting Rate Model |  |  | Optimal Linear Estimator |  |  |
| --- | --- | --- | --- | --- | --- | --- | --- | --- |
|  | Scenario | n | MTE | 95% CI | PE-2020 | MTE | 95% CI | PE-2020 |
| a. Tasmania-wide |  |  |  |  |  |  |  |  |
|  | Physical specimens only (P) | 99 | 1937 | 1934–1938 | 1 | 1939 | 1935–1940 | 1 |
|  | P + unconfirmed kills/captures | 162 | 1975 | 1953–1999 | 1 | 1986 | 1957–2019 | 0.975 |
|  | P + multiple sighting witnesses | 198 | 1960 | 1935–1995 | 1 | 1971 | 1936–2023 | 0.947 |
|  | P + only highest-rated sightings | 312 | 1987 | 1958–2003 | 1 | 1999 | 1962–2024 | 0.928 |
|  | P + uncertain until 1950 cut-off | 376 | 1951 | 1948–1951 | 1 | 1951 | 1948–1952 | 1 |
|  | P + all ‘expert’ sightings (E) | 429 | 2002 | 1986–2015 | 1 | 2006 | 1987–2023 | 0.806 |
|  | All records (default weights) | 1237 | 2011 | 1997–2019 | 0.999 | 2016 | 1999–2026 | 0.598 |
|  | All records (low weights) | 1237 | 1994 | 1970–2018 | 0.986 | 2007 | 1975–2040 | 0.834 |
| b. Bioregional (P + E records) |  |  |  |  |  |  |  |  |
|  | South-East | 24 | 1972 | 1920–1997 | 1 | 2012 | 1925–2132 | 0.657 |
|  | North-East | 68 | 1985 | 1970–1996 | 1 | 1993 | 1971–2018 | 0.980 |
|  | North-West | 167 | 1991 | 1972–1998 | 1 | 2001 | 1977–2015 | 0.999 |
|  | West / World Heritage Area | 264 | 1999 | 1978–2015 | 1 | 2008 | 1983–2031 | 0.794 |

*Footnotes.* **a.** Assessment using an aggregate of records from across the island, from 1910–2019, based on various combinations of verified physical specimens only, physical records and the addition of unconfirmed kills and captures, or all unverified expert sightings, or all record types (including physical records, expert sightings, and other observations by the public), weighted by the record’s quality rating using default or low assigned probabilities. **b.** Spatially disaggregated scenarios, using records specific to four main bioregions, based on physical specimens and expert sightings only. Shown for each scenario is the number of records sampled (n), mean time of extinction (MTE, calendar year) and the 95% confidence intervals of the simulations (95% CI, based on distribution percentiles), and the modelled probability of extinction in the year 2020 (PE-2020) (Brook et al. 2019). The two alternative statistical extinction-date estimators, used for inference of extinction date based on sightings records, are the variable sighting rate model of McInerny et al. (2006) and the optimal linear estimator of Roberts & Solow (2003). The individual IBRA7 Bioregions included within each geographic agglomeration of sighting records are as follows: South-East = Northern Midlands, South East; North-East = Ben Lomond, Furneaux; North-West = Northern Slopes, King; West / World Heritage Area = Central Highlands, West, Southern Ranges.

Relaxing the constraints further, if all unverified expert reports are considered by down-weighting them proportionally according to their quality rating (as per Brook et al. 2019)—the projected extinction window spans a later period, from the 1980s to the present, although the probability of ongoing persistence to 2020 is low (Table 2a; Fig. 1a,b). This might seem surprisingly recent, but is supported by, for example, the concerted official search efforts from Parks & Wildlife authorities in the 1980s that were motivated by apparently highly credible sightings (Smith 1981). Finally, in the most liberal treatment of the sighting record, where all records, including opportunistic sightings by the public, are used, yields a similar conclusion. If low inclusion probabilities are assigned to each type of observation, then the extinction window spans 1970–2018 for the variable sighting rate model, with a median estimated year of 1994.

A sensitivity analysis shows our results to be robust to permutations in the record-inclusion criteria or assignment of sighting probabilities (Fig. 1c and Appendix S4).

### Pattern of extirpations

The Tasmania-wide analysis (Table 2a) combines sighting records of the Thylacine from across the island, irrespective of location. However, extinction often progresses via an intermediate process of range contractions and spatially heterogenous declines, themselves driven by a variable local intensity of threats like habitat change and hunting. A corresponding analysis of bioregional clusters of sightings, based on EDEs fitted to the physical-specimen and expert-sightings data (Table 2b), reveals a general pattern of local losses starting in south-east and midland regions of the island (abutting areas where grazing, agriculture and settlement was concentrated) and later extending to the remote wilderness areas of the center and south-west (see also Table 1, where the last physical records indicate final refuges in the west and south). This results in a median extinction date of 1999 or 2008 (for the variable sighting rate and OLE models, respectively), but the confidence intervals for these extirpations are wide for many regional aggregations, with two from the OLE even overlapping with the present (Table 2b).

For greater spatial fidelity and a visualization of the dynamical time course of range decline for the Thylacine, the results are most usefully mapped as a geographical projection of point-wise extinction-date estimates on a 0.1° latitude-longitude grid, with the contribution of sighting records surrounding each landscape-grid point down-weighted using a distance-decay function, while retaining each record’s respective sighting probability. This results in the multi-weighted contour surfaces illustrated in Fig. 2, superimposed on a coastal outline of Tasmania.

**Fig. 2.**
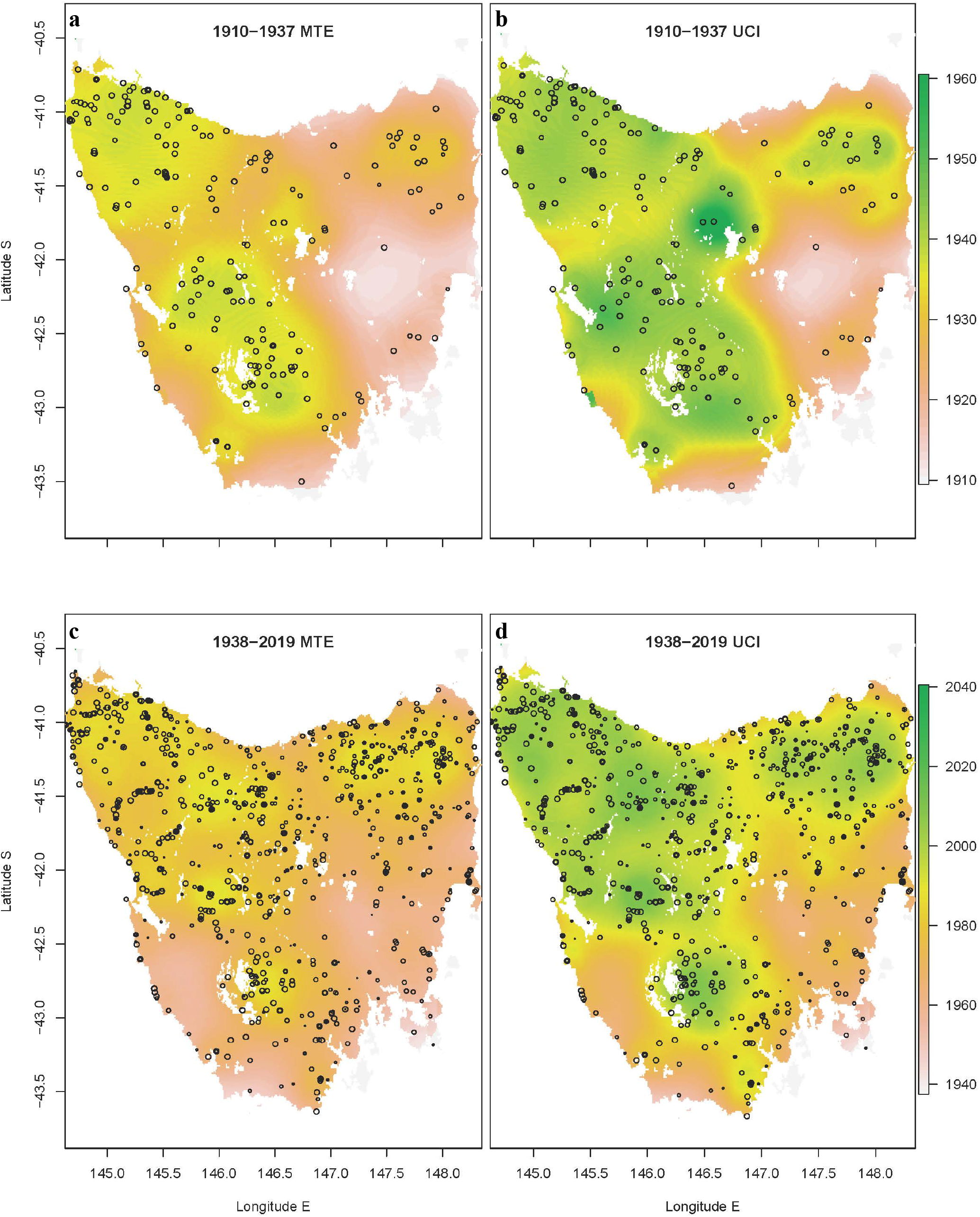
Spatial extirpation pattern for the Thylacine in Tasmania. Colored contour maps of the inferred year of local extirpation, estimated for each pixel across a 0.1° geographical grid of the island (area = 64,519 km^2^). The results were generated by fitting a re-sampled (Brook et al. 2019) variable sighting rate extinction-date estimator (McInerny et al. 2006) to the observational record, with sightings down-weighted by their assigned uncertainty and the great-circle distance from the target grid cell (see Methods). **a**. mean time of extinction (MTE) and **b**. upper confidence interval (UCI) for records spanning 1910–1937 (a mix of verified and uncertain records, n = 258). **c.**, **d.**, as for a., b., except using only records from 1938 onwards (all uncertain records, n = 979). The circles in each plot show individual sightings, sized by rated quality: 5 (highest quality) for the largest circles, down to 1 (lowest) for the smallest.

The situation for the species by 1937 (based on a mixture of records on kills, captures and expert sightings) was of a severe decline across most of the landscape, with pockets of probable persistence in south-central and north-west regions of the island, likely connected by dispersal corridors (Fig. 2a,b). There was also some possibility of a remnant isolated sub-population persisting in the north-east at that time, but with strong evidence for an early extirpation (by the 1920s) in the midlands and along the south-east coastal region, where the bounty killing had been particularly intensive (Prowse et al. 2013). From 1938 onwards, all records (through to 2019) are of unverified sightings or reported (but not confirmed) kills and captures, of varying quality, coming from experienced trappers through to the (largely) Thylacine-naïve public. These data (Fig. 2c,d) indicate extirpation by the early 1960s across most of the southern half of the state, with longer-term persistence along a band stretching across the Tasmanian Wilderness World Heritage Area, from Lake Pedder in the south-center across to the western edge of the central highlands and up to the Tarkine in the north-west. There is also remarkable congruency across the two sets of extirpation maps in the geographic location of potential refugia (‘hotspots’ of records), despite being based on assessment of completely temporally separate, non-overlapping data (the latter period has no certain, verified records). The most likely termination date for the species seems to have occurred within this zone by the late 1990s, although the upper confidence bounds of the model include the present day in some wilderness regions of the island.

## Discussion

Based on the extensive (albeit uncertain) sighting record that post-dates the death of the last captive Thylacine in 1936—which we integrated using spatially explicit inferential methods that account for mixed-quality observations—it seems reasonable to conclude that a remnant population of the species persisted in remote areas of Tasmania for many years thereafter. This assertion has been argued on other ecological and detectability grounds (Guiler & Godard 1998; Bulte et al. 2003; Terry 2005; Lang 2014; Sleightholme & Campbell 2016), however ours is the most comprehensive assimilation, model-based quantification, and spatio-temporal visualization of the full body of Thylacine sighting data. But why then, if the species persisted for decades after the 1930s, did the supply of live specimens and carcasses cease and, moreover, why was the species never photographed in the wild or confirmed by other scientific field methods?

The fate of the last wild individual of a species is rarely witnessed by people. This is especially the case for species like the Thylacine, which ranged widely but sparsely across large swathes of the Tasmanian wilderness (Sleightholme & Campbell 2016). The last survivors were probably increasingly difficult to detect as they became ever more wary of potentially fatal interactions with people, as numbers dwindled, and as the species’ spatial distribution contracted and disaggregated as the species’ population decline progressed (Guiler & Godard 1998; Fisher & Blomberg 2011). A direct reading of the physical evidence implies extinction in the wild by the mid-1930s. However, when species are driven to extreme rarity, most of the final records will be uncertain/unverified sightings; in the case of the Thylacine, after formal legal protection in 1936, there was a disincentive to self-report kills, for fear of penalty or prosecution (Brook et al. 2018).

Regarding verified detections, modern remotely triggered instruments are among the most cost-effective, unobtrusive and failsafe ways to record elusive vertebrate wildlife, and well-disguised cameras have been used to rediscover cautious carnivores that were previously thought extinct, such as the Zanzibar leopard (*Panthera pardus adersi*) (Goldman & Walsh 2002; Li 2018). However, digital-trail-camera technology has only been widely deployed in field ecology over the past two decades (Meek et al. 2015), with earlier visual searches and film-camera field operations being of relatively short duration and restricted geographical coverage (Guiler 1966; Griffith 1972; Smith 1981). Similarly, from the public’s perspective, it has only been during this recent time span that smartphones and vehicle dashboard cameras have been in widespread use.

Although the extinction-range estimates reported herein span a more recent period than the estimate (1936–1943) derived previously from a Bayesian model of mixed-certainty sightings (Carlson et al. 2018), we have criticized the latter for only using a small fraction (<10%) of all possible records (Brook et al. 2018). The Bayesian approach, when used on annually aggregated data, is insensitive to information embodied in the uncertain sightings, at least for the Thylacine, with only 1-3 years difference in the extinction-date estimate derived after rejecting all uncertain records, compared to modelling certain and uncertain records separately (Brook et al. 2018).

The most vexing difficulty with a scientific mystery like this—trying to decide which sightings are correct, and which are false—is quantifying the risk of ascertainment bias. During the post-bounty period, Thylacine encounters were noted as being rare, but because there were still occasional kills and captures, the species was known with certainty to persist. During this time, unverified sightings were made regularly (128 reports between 1910 and 1936), and there seems little reason to doubt their authenticity, there being no general perception at the time that an unproven sighting was anything particularly remarkable (Sleightholme & Campbell 2016). However, in the years following the death of the last captive Thylacine, when zoos sought new specimens (offering substantial remuneration) and yet none could be secured, interest in proving the species’ ongoing existence steadily rose, such that by the 1960s it was recognized as a puzzling quandary (Guiler 1966; Griffith 1972). As recognition of this evidence gap grew, there was a greater incentive to falsely report sightings (for notoriety), or even a subconscious desire to *want* to see a live Thylacine, leading to inflated misidentification errors. Without the reassurance of an occasional ‘ironclad’ (physical) record, the time point at which there was a switch from some sightings being true, to *all* being wrong (i.e., after extinction), is left inevitably shrouded in the conservation-biology equivalent of a ‘fog of war’. To try and cut through this, we are left with an inescapable reliance on observer credibility and the associated sighting details (much as is done when weighing up the usefulness of eyewitness testimonies in law courts; Wechsler et al. 2015), the specifics of which are listed for each record in the multivariate meta-data of the TTSRD. An extended discussion of this topic is given in Appendix S2 (Supporting Information).

Regardless of which scenarios or assumptions one chooses to favor or discount, our new method for mapping spatial contours of extirpation dates is useful not only for reconstructing the dynamics of the Thylacine’s range contraction, but also for identifying the most likely spatial refugia occupied by the species prior to its range-wide extinction. Indeed, the extirpation-mapping algorithm we developed for the Thylacine case study is general and could be applied equally to other species of conservation concern that are verified to survive but where the synthesis of sighting records (confirmed or uncertain) has previously defied integration. Additionally, these extirpation-probability maps could be unified with existing habitat- and climate-envelope methods (using occurrences recorded prior to a species’ decline), to pinpoint regions where both available niche space and recent sightings indicate localities of potential survival, as a means of targeting more intensified search efforts, restoration, rewilding, or reallocation of resources when persistence is extremely unlikely (Rout et al. 2010).

In conclusion, this collective body of evidence and associated analyses indicates that while the Thylacine is unlikely to persist to the present, the true extinction year likely occurred much later than the commonly held date of 1936, when the last captive animal died. Indeed, our scenario analysis on new sighting-record compilation implies that the inferred ‘extinction window’ is wider and more recent than suggested by previous modelling (Carlson et al. 2018), spanning from the 1960s to the present day, with the peak likelihood centered on the late 1980s to early 2000s. Certainly, these aggregate data and modelling point to the species persisting within the remote wilderness of the southwest and central highlands regions of island for decades after the last confirmed specimen. Finally, we note that although our findings for this iconic species hold intrinsic value, our new spatio-temporal mapping of extirpation patterns is also applicable more generally, to support the conservation prioritization and search efforts for other rare taxa of uncertain status (Solow 1993; Jarić & Roberts 2014; Solow & Beet 2014).

## Supporting information

TTSRD

## Acknowledgments

The authors are indebted to C. Bailey, T. Gordon (Queen Victoria Museum and Art Gallery, Launceston), R. Smith (QVMAG) and R. Gaffney (DPIPWE, Hobart) for their assistance in sourcing sightings records and providing records from their own archival material, and M. Lee for a useful critical discussion on ascertainment bias.

## Supporting Information

Extended discussion on the historical context of thylacine sightings (Appendix S1) and a critical analysis on what the TTSRD compilation can (and cannot) tell us about the Thylacine’s extinction, given the limits of knowledge and inference (Appendix S2). Supporting references are listed in Appendix S3, and Appendix S4 provides Tables S1-S7, giving detailed breakdowns of the TTSRD records by type, quality, sighting class, probability assignments, and regional scenario analysis for specific EDE models, and Fig. S1-S2, which show the time-series of Thylacine sightings, disaggregated by observer type and quality rating.

## Data and materials availability

Data analysis and modelling were implemented in Program R v.4.0.2 (http://r-project.org). The source instruction code, functions and input data files required to reproduce all results reported herein are publicly available on GitHub (http://github.com/bwbrook), along with The Tasmanian Thylacine Sighting Records Database (TTSRD) as CSV file or a Microsoft Excel workbook, and images (attachments).

