## Supporting Information for "Resolving when (and where) the Thylacine went extinct"

#### Appendix S1

##### Historical Context of Thylacine Sightings

The Thylacine was the largest living predatory marsupial following the extinction of most of the Sahul mammalian megafauna in the late Pleistocene (Prowse et al. 2014). Much debate remains over exactly when the extinction occurred. Was it when the last individual died in the Hobart zoo in 1936? Or was it at some later unobserved date? Or does the species persist through to the present day, yet remains cryptic and extraordinarily difficult to find?

In Tasmania, the Thylacine was known from the earliest days of settlement, although it was reported to be elusive (Jeffreys 1820). The original Tasmanians had coexisted with the Thylacine for over forty millennia prior and undertook limited hunting of the species (Guiler 1985). Following European colonisation starting in 1803, the native Tasmanians were displaced from the landscape by disease, persecution, and war, and in 1833 the last few hundred indigenous people were 'relocated' to Flinders Island. Meanwhile, in 1886 the Tasmanian government yielded to pressure from a small but vocal section of pastoralists and put a price on the Thylacine's head, paying £1 per adult and 10 shillings per young. After some delays, the first bounty was paid out in 1888 and the scheme ran until its termination in 1908, although payments continued to be made into 1909. The Van Diemen's Land Company's also ran a bounty scheme from 1830 until its termination in 1914, with a two-year hiatus from 1838–1840. A number of local bounties ran in concert.

By 1910, Thylacines were so rare that zoos offered substantial rewards for live specimens (Guiler 1985). Encounters in the wild occurred sporadically across much of Tasmania over the next few decades, but interactions between Thylacines and people were increasingly uncommon. Of the few animals subsequently captured, most were sent to Australian zoos or sold to overseas collections. Approximately 125 Thylacines were displayed in thirteen zoos from 1900 to 1936. Despite the relatively large monetary sums offered for their capture, few Thylacines were caught for the zoo trade after 1925. In 1930 the last confirmed shooting of a wild Thylacine occurred near Mawbanna (TTSRD record ID #109), and on 7th September 1936, the last absolutely verified live Thylacine—a male captured over five years earlier (#127)—died at the Hobart Zoo (Sleightholme et al. 2020). And so, this unique, wolf-like carnivorous marsupial had been driven to near extinction by human persecution, after which small population processes apparently dealt the final blow. However, some researchers claim, through modelling and documentary analysis, that bounty hunting alone is insufficient to explain the Thylacine's demise, invoking disease or habitat loss as additional causes (Bulte et al. 2003; Paddle 2012; Haygarth 2017).

Given this context, it is highly probable that the species persisted beyond the widely cited 1936 extinction date. The two leading authorities on the species (Guiler and Möeller) were also of this opinion. Even Nick Mooney, the Parks & Wildlife officer who was for decades responsible for collating and investigating Thylacine sightings, is on record as stating (in 2013) that the species persisted beyond the death of the last captive specimen:

"*As to the fate of those last 100 thylacines... my suggestion is they were fragmented into small groups, some of which just fizzled out like flags being brushed off a war game*" (quoted in Bevilacqua 2013).

The sighting record and mean-time-of-extinction map (main paper, Figure 2a) closely reflect the distribution map in Sleightholme & Campbell (Sleightholme & Campbell 2016). At that point in time, the Thylacine seems to have been functionally extinct to the east of a line running from Launceston to Hobart. This is supported by documentary evidence from the time. In the 1920s and 1930s, Arthur Reid, the curator of Hobart Zoo, travelled the length of the east coast of Tasmania, trying to obtain Thylacines for the zoo, but was unable to do so. All the trappers he spoke to reported that they had not seen any Thylacines in years. *The Mercury* of the 15 May 1923 (p.8) reported on a meeting of the Royal Society of Tasmania in which Arthur Reid gave an account of a recent journey along the east coast of Tasmania to obtain animals for the zoo:

*"The whole journey, he said, occupied about 400 miles, and he went in and out of all the ports and townships as far as Musselroe, on the East Coast. He had been told that he would find enormous numbers of forester kangaroo, but he was very disgusted with the search, as they failed to bring home one. He saw very few wallaby, opossum, or kangaroo. During the whole of his travel, Mr Reid said he failed to see one Tasmanian marsupial wolf, which species, he said, must be extinct on the East Coast, and as they have not one at the Zoo, their only possibility of obtaining one was on the West Coast".*

Reid's comments are significant in that, after a journey of 400 miles (640 km), taking in all the east coast ports and towns, he neither observed a Thylacine, nor was offered the opportunity to purchase a specimen.

With reference to the possibility of a refuge population around Ben Lomond in the North East, Sleightholme & Campbell (Sleightholme & Campbell 2016) state:

*"It now seems virtually certain that the thylacine became extinct, or functionally so, in the developed, eastern half of the state during the early 1920s. That said, there is some evidence*

*to suggest that a relict population, probably numbering no more than a few individuals, survived in the area to the north of Ben Lomond into the early 1930s (Collins, [Mt. Direction, 1930], Lawes, [Cuckoo Valley near Mt. Stronach, 1930], Woods, [Ringarooma, 1934-5], and Peck, [Mt. Barrow, 1935-6])".*

It is worth noting that dating discrepancies and misidentifications may account for most of these alleged sightings, as all the reports were made retrospectively, many years after the sightings took place. For example, Collins dates his sighting with his recollection of seeing a tiger at the old Beaumaris Zoo: i.e., he states (see #117, attachment) "*around this time*", and gives the date as 1930, but his sighting was undoubtably earlier, as the zoo had moved to its new location on the Domain in 1921. Woods recalled seeing the eye reflection of what he thought was a tiger illuminated by his flashlight when inspecting snare lines at night. This sighting might be a simple case of misidentification. The fact that no kills or captures were recorded in the Ben Lomond area during the 1930s further weakens the north-eastern survival argument. That said, there are still wild corridors of bushland linking the central and south western parts of the state to the north east, so it is possible that migration of the relict population occurred along these natural corridors to account for later sightings.

Parks & Wildlife Tasmania certainly took the Mooney investigation (launched following the well-known Naarding sighting, #401) seriously, and stated so in their official report:

"*It was concluded that the search area was used at least irregularly by Thylacines up until Autumn 1982, but use had diminished due to increased disturbance to the point that detection of animals is not probable, despite large efforts. Unless the Thylacine observed in March 1982 by the service biologist was the last of the species, it must be accepted that Thylacines survive in a number of areas in Tasmania*".

With respect to the conclusions drawn by the search undertaken by Griffith, Malley & Brown in the late 1960s and early 1970s (Griffith 1972), they state:

*"Having been into every likely corner of Tasmania, we appreciate how elusive a few Thylacines could be in that country. That we haven't found conclusive evidence of Thylacines does not necessarily indicate that they are extinct. Rather, it illustrates the enormity of our task. We are powerless to help Thylacines unless we can find them and assess their predicament: the alternative is that an uncertain case of extinction will become certain".*

One of the authors (SS) knows, from sight of his private correspondence with Professor Heinz Möeller, that Dr. Eric Guiler from the University of Tasmania—the leading expert on the species in the second half of the 20$^{th}$ Century—was convinced that Thylacines still existed in the Arthur River area in 2002. Indeed, he was investigating a spate of recent sightings from this vicinity when he suffered his near fatal stroke. Further, the 1995-1997 Clarke related sightings, two of which [#937 & #1294] were rated by Parks & Wildlife as "*the most promising sightings in 10 to 20 years*", were located geographically in exactly the region where the current study (Fig. 2c,d) shows they should in the late 1990s. Even the National Museum of Australia (2019) states: *"Thylacines are functionally extinct which means that any remaining population is so small that the species no longer plays a significant role in the ecosystem".* Consequently, doubt exists.

Historically, the Thylacine ranged across Tasmania from the east to the west coasts. Guiler & Godard (Guiler & Godard 1998) estimated that at the time of colonial settlement in 1803, the total Thylacine population would have numbered between 2000–4000 individuals. Concerns over the sudden decline in the Thylacine population at the beginning of the 20th century were expressed by many leading zoologists and naturalists of the day: Le Souëf in 1907, Roberts in 1909, Flynn in 1914, Lord in 1919, Pearson in 1936, Fleay in 1937 & Sharland in 1938, leading to many searches and expeditions: Summers (1937), Fleming (1937), Fleming / Sharland (1938), Fleay (1945/1946), Guiler (1959; 1960; 1961; 1963-64 & 1980-81), Griffith, Malley & Brown (1968-1972), Smith & Pyrke (1978-1980), Mooney (1982-1983) & Wright (1984). Many reported sightings in the 1940s and 1950s were made by multiple persons sighting the same animal, or by bushmen or farmers with first-hand knowledge of the Thylacine. Details of all of these dated expeditions cited above are provided in past publications on the Thylacine (Guiler 1985; Owen 2003; Sleightholme & Campbell 2016)

#### Appendix S2

##### What can the TTSRD compilation tell us about the Thylacine's extinction?

A clear-eyed appraisal of the TTSRD records reveals no obvious qualitative difference between sightings made before and after 1936, when the last captive individual died. It is not clear, *prima facie*, that all post-1936 sightings can be summarily dismissed, given that many of the accounts (rated highly by authorities, and us in the TTSRD) hold up to careful scrutiny. With the compiled records of the TTSRD, we are in a unique position in being able to make comment—and for the first time undertake a rigorous analysis—on the extinction question.

The most conservative approach to considering the Thylacine sighting record is to only evaluate confirmed records and reject all others. If we do this, the conclusion would be that the species last stronghold was in the north west of the state and that the final date of extinction was in the early 1930s. Less strictly, all records post-dating the last kill or capture must be considered possible, but uncertain. Logic says that the more time that has passed since the last confirmed record, the less likely *any* record is of being correct. Conversely, the most overtly liberal approach to these data is to assume that all sightings are valid. This leads to the naive conclusion that the species *must* continue to persist in remote Tasmania. Reality surely lies somewhere in between these two extremes. The question then arises, how best to treat these (majority of) uncertain records? Should this information be discarded on the basis of it not being absolutely verifiable, or can it be used probabilistically to make a stronger inference about the most likely date of extinction, or indeed whether there is any reasonable likelihood that the species continues to persist to the present day?

First, to the primary data. Tables S1 and S2 give a breakdown of sightings from 1910 onwards, broken down by type, class, and quality rating. Of the records with sufficient detail given, 376 were from along a road (96.8% of these were made from a vehicle), and 779 observations were reported by people undertaking activities within the bush (off-track, in the scrub). Most were made by a lone observer (66.1%) of single animals (85.0%) during the day (60.8%) at an estimated distance of less than 30 metres (85.0%). Yet a remarkable 24 sightings were made by groups of three or more people, which are difficult (albeit not impossible) to explain away as a group delusion.

Why then, has no scientifically verifiable evidence been presented on the Thylacine since the 1930s? In fact, unequivocal observations of numerous species ceased after their discovery (and apparent extinction date), often for decades or centuries until their eventual re-discovery

(Scheffers et al. 2011). For example, there was no verifiable evidence of the Leadbeater's possum (*Gymnobelideus leadbeateri*) for 94 years after it was believed to have become extinct. Similarly, there were no confirmed sightings of the bridled nail-tail wallaby (*Onychogalea fraenata*) between 1937 and 1973. The Takahe (*Porphyrio hochstetteri*) of New Zealand was considered extinct by 1898 and was only rediscovered in 1948. The Bermuda Petrel (*Pterodroma cahow*) was thought to have become extinct in the 1620s, but was rediscovered in 1951, over 300 years later. Other cases are detailed—along with an accompanying EDE analysis that accounts for certain and uncertain sightings—in Brook et al. (2019). These examples clearly demonstrate that species can persist without verifiable evidence, especially if their habitat has not been intensively surveyed. As Baumsteiger & Moyle (2017) note:

"*Determination of extinction can be surprisingly complicated. It is often difficult to say with certainty when the last individual is gone because most lineages are cryptic at small population sizes, making it difficult to determine 'no reasonable doubt'*".

Although Tasmania is geographically restricted, the island is large (68,401 km$^2$) and much of the west, central and south-west regions are lightly populated or protected wilderness. Previously, in response to a Bayesian model of extinction (Carlson et al. 2018), we have offered an explanation as to why the 'supply' of physical specimens or live Thylacines in Tasmania might have ceased many years, or even decades, prior to its eventual extinction (Brook et al. 2018).

From the perspective of modelling the TTSRD, one is obliged to form a default viewpoint and that determines the most appropriate EDE (extinction date estimator) model to use:

(i) The null hypothesis that the species persists, unless proven extinct (which is typically impossible, at least for cryptic species occupying large landscapes). In this case, the question the modelling must ask is: "**Given the sighting record, what is the (statistical) likelihood of failing to secure any *absolute proof image or capture* since 1931?**". In this case, one has no basis for assigning any sighting a probability of zero, although one could, in theory speculate (and model) trends (or even step changes) in these probabilities over time. The assumption of fixed sighting probabilities per class remains open to debate, but difficult to resolve. In this case, an inference method like that used in the main paper (Brook et al. 2019), which treats all observations probabilistically using a frequentist framework, will be most appropriate.

(ii) As an alternative hypothesis, one could assume that the species is extinct. The question then becomes: "**Given that the species went extinct but at some unknown date, what is the**

**probability that any given sighting is correct?**". The question of interest here involves the time series of uncertain records after the last verified record. In the unknown period when the species was extant but unverified, the likelihood that a given observation was true will be some probability between 0 and 1. After extinction, it will be precisely 0. In this case, a Bayesian method like that of Solow & Beet (2014) *might* be more informative, although as noted earlier, it has a strong tendency to upweight certain over uncertain records, as was the case for a previous modelling on annual occurrence data (Brook et al. 2018; Carlson et al. 2018).

A philosophical exploration of the advantages and shortcomings of applying frequentist versus Bayesian paradigms for inferring extinction dates given a sighting record of certain and uncertain records, is given in the Supplementary Information of Brook et al. (2019).

Further, under either viewpoint (where we define E = expert sightings and O = other observations), we might hypothesize that the E-type sightings would decline over time even with no change to the Thylacine population, because those considered most reliable were verified eyewitnesses of encounters with Thylacines in the early decades of the 20$^{th}$ century; these people have aged and died over subsequent decades, and their memories faded. By contrast, we might expect O-type sightings to increase proportionally over time, because fewer observers closer to the present, other than wildlife scientists, can ever be considered experts, and as time passes, there is increasing notoriety in making a 'sighting'. For the alternative hypothesis (ii, above), for both E and O, there will be some background error rate (that might be relatively time invariant, before and after extinction) and some true sighting rate (which is positive if the species persists, and zero after it goes extinct).

From a logical standpoint, what can be made of the high-quality sighting data that have continued to appear sporadically, where the witness (or witnesses) appears reliable and/or knowledgeable about the species, and stands to gain nothing from making the report? These include sightings made by bushmen (including fur trappers), farmers, forestry staff and police officers, all of which are unlikely to have made an error in identification. It is difficult, on the face of it, to exclude any of these *a priori*. A number of possible interpretations seem valid:

1) Extreme pessimism: the post-1936 reports were all wrong/false (i.e., all sightings were actually some combination of dogs, devils, quolls, cats, illusions or tall tales);

2) Extreme optimism: the reports were all correct (in this case we simply assess the full record without any attached probabilities and project onwards from the last sighting);

3) Ambivalence: some reports were correct, some mistakes (in this case, one needs to attach uncertainty to each record, based on the observer, the circumstances, and the overall quality of the report, as we have done in the analysis that accompanies the main paper);

4) Temporal ambivalence, then certainty after an uncertain date: the reports were a mix of correct and false sightings up until some undetermined time point, after which the species actually went extinct; thereafter, *all* subsequent records were, by definition, false.

In case 1, the interpretative task is straightforward. The time of extinction is assessed based only on the physical records, yielding a best-estimate date of 1935–1939 (see Table S8). This conclusion is insensitive to the EDE method used for inference. A Bayesian analysis using only a single record per year, graded as either certain or uncertain, yields a similar answer (Carlson et al. 2018) with the consequence of effectively down-weighting uncertain sightings to near zero (Brook et al. 2018).

In case 2, the latest TTSRD record graded as quality 4 or 5 (see Tables S1 and S4) was in 2018 (an O4, #1397), or 2014 (a S4, #429). Under this line of argument, the high-quality sighting record can be taken as a direct insight into persistence, with no need for an EDE. The conclusion, under this assumption, is that the species was extant within the last decade.

Case 3 is the one upon which we have focused analysis and discussion in this paper. The working hypothesis here is that the species remains extant (null hypothesis), and then 're-sampling with EDE methods can be used to infer the likelihood of a temporal gap of any given length after the last accepted record (itself stochastic), given that the null hypothesis is true. We use this approach to construct confidence intervals, and so to reject persistence beyond a given year (defined by the probabilities generated by the EDE model with uncertain records).

Case 4 is the most difficult to handle. One possibility is to seek to detect some pattern in the sightings records, so as to identify a switch point quantitatively, when the observational record went from being composed of a mix of correct and false sightings, to one in which all sightings were axiomatically false (after extinction). One approach is to undertake a 'stress test' of the EDE-with-uncertainty analysis, using extremely high rejection probabilities for individual sightings, such as the P(L) scenario shown in Table S3 and incorporated within Fig. 1c. Another is to include or exclude certain sighting types from the analysis and assess the impact on the inferred extinction year. This is shown in Table S5 (contrasted to Table 1b), and with bioregional breakdowns shown in Tables S6 and S7. The upshot of these analyses is that the conclusion of late survival of the Thylacine—to the end of the 20$^{th}$ Century or beginning of the 21$^{st}$ Century—is robust to the quality of the sighting records that are used.

Time-series plots (Fig. S1 and S2) show a rise then fall in expert high-quality sightings over time, a similar but lagged response for non-expert high-quality sightings, and a peak then a relatively constant rate for lower-quality sightings. Our interpretation of this pattern is that extinction occurred sometime after 1985, when the frequencies decline for all types. Further to this point, within the TTSRD, there are 32 high-quality (S4/S5) sightings for the 1960s, 20 for the 1970s, 12 for the 1980s, 8 for the 1990s and 1 for the 2000s. Setting a probable extinction date too early (e.g., the 1960s) based on an arbitrary decision about when to reject all dates amounts to discounting many sightings made by experienced bushmen (Blacklow, Pursell, Summers, Blizzard, Blackwell, Williams, King, Porteus), as well as Naarding's 1982 sighting, and the Roberts sighting (#1374) which Mooney described as being "very credible".

In sum, this collective body of evidence and associated analyses indicates that the continued persistence of the Thylacine is unlikely—but possible—and that the true extinction year, if the species is indeed now extinct, occurred much later than the commonly held date of 1936.

#### Appendix S3

#### Supporting Information References


Baumsteiger J, Moyle PB. 2017. Assessing extinction. BioScience **67**:357-366. DOI: 10.1093/biosci/bix001

Bevilacqua S. 2013. Dreamers flesh out hopes that the thylacine still exists in Tasmania's wilderness.

Brook BW, Buettel JC, Jarić I. 2019. A fast re-sampling method for using reliability ratings of sightings with extinction date estimators. Ecology **100**:e02787. DOI: 10.1002/ecy.2787

Brook BW, Sleightholme SR, Campbell CR, Buettel JC. 2018. Deficiencies in estimating the extinction date of the thylacine with mixed certainty data. Conservation Biology **32**:1195-1197. DOI: 10.1111/cobi.13186

Bulte EH, Horan RD, Shogren JF. 2003. Is the Tasmanian tiger extinct? A biological–economic re-evaluation. Ecological Economics **45**:271-279. DOI: 10.1016/s0921-8009(03)00076-4

Carlson CJ, Bond AL, Burgio KR. 2018. Estimating the extinction date of the thylacine with mixed certainty data. Conservation Biology **32**:477-483. DOI: 10.1111/cobi.13037

Griffith J. 1972. Search for the Tasmanian tiger. Natural History **81**:70.

Guiler ER 1985. Thylacine: The Tragedy of The Tasmanian Tiger. Oxford University Press, Melbourne, Australia.

Guiler ER, Godard P 1998. Tasmanian Tiger: A Lesson To Be Learnt. Abrolhos Publishing, Perth, Western Australia.

Haygarth N. 2017. The myth of the dedicated thylacine hunter: stockman–hunter culture and the decline of the thylacine (Tasmanian tiger) in Tasmania during the late nineteenth and early twentieth centuries. Tasmanian Historical Research Association **64**:30-45.

Jeffreys CH 1820. Geographical and descriptive delineations of the island of Van Diemen’s Land. J.M. Richardson, London.

Owen D 2003. Thylacine: The Tragic Tale of the Tasmanian Tiger. Allen & Unwin, Sydney, Australia.

Paddle R. 2012. The thylacine's last straw: epidemic disease in a recent mammalian extinction. Australian Zoologist **36**:75-92.

Prowse TAA, Johnson CN, Bradshaw CJA, Brook BW. 2014. An ecological regime shift resulting from disrupted predator-prey interactions in Holocene Australia. Ecology **95**:693-702. DOI: 10.1890/13-0746.1

Scheffers BR, Yong DL, Harris JBC, Giam XL, Sodhi NS. 2011. The world's rediscovered species: back from the brink? PLoS One **6**. DOI: 10.1371/journal.pone.0022531

Sleightholme SR, Campbell CR. 2016. A retrospective assessment of 20th century thylacine populations. Australian Zoologist **38**:102-129. DOI: 10.7882/AZ.2015.023

Sleightholme SR, Gordon TJ, Campbell CR. 2020. The Kaine capture - questioning the history of the last Thylacine in captivity. Australian Zoologist **41**:1-11. DOI: 10.7882/AZ.2019.032

Solow AR, Beet AR. 2014. On uncertain sightings and inference about extinction. Conservation Biology **28**:1119-1123. DOI: 10.1111/cobi.12309

### Appendix S4

#### Supporting Tables and Figures

**Table S1.** Frequency table of TTSRD records by type and quality. Physical and expert records were never rated as <3 quality.

| | | | *Quality Rating* | | | |
|---|---|---|---|---|---|---|
| *Type* | 5 | 4 | 3 | 2 | 1 | Description |
| PS | 85 | 8 | 6 | NA | NA | Physical specimen (kill or capture, confirmed) |
| EO | 227 | 77 | 34 | NA | NA | Expert observation (sighting, or kill/capture with no confirmed specimen) |
| EI | 71 | 14 | 6 | NA | NA | Expert index (track, vocalisation, predation, den, etc.) |
| OO | 76 | 210 | 135 | 121 | 59 | Other observation (sighting, plus a few dubious claimed kills/captures) |
| OI | 2 | 19 | 22 | 54 | 11 | Other index (track, vocalisation, predation, den, etc.) |

**Table S2.** Frequency table of TTSRD records by sighting class.

| Sighting Class | # |
|---|---|
| Capture - confirmed (C) | 66 |
| Kill - confirmed (K) | 33 |
| Capture - unverified (C*) | 28 |
| Kill - unverified (K*) | 35 |
| Expert visual sighting (S) | 282 |
| Expert indirect record (SI) | 91 |
| Other visual sighting (O) | 427 |
| Other indirect record (OI) | 43 |
| Other low-quality visual (O*) | 167 |
| Other low-quality indirect (O*I) | 65 |

**Table S3.** Relative probability (P) weightings of records by type (PS = physical specimen, EO = expert observation, EI = Expert indicator of presence, OO = other observation, OI = other indicator), for the low (L), default (D) and high (H) scenarios.

| Type | P(L) | P(D) | P(H) |
|---|---|---|---|
| PS | 1 | 1 | 1 |
| EO | 0.05 | 0.25 | 0.5 |
| EI | 0.01 | 0.1 | 0.2 |
| OO | 0.005 | 0.05 | 0 |
| OI | 0.001 | 0.01 | 0 |

**Table S4.** Probability multipliers ($P_{mult}$) on the record weightings given in Table S3, depending on the quality rating (QR) of the observation. For example, a record of QR = 5 (highest quality) is unaltered, whereas a QR = 1 has only 20 % of the weighting.

| QR | $P_{mult}$ |
|---|---|
| 5 | 1 |
| 4 | 0.8 |
| 3 | 0.6 |
| 2 | 0.4 |
| 1 | 0.2 |

**Table S5**. Scenarios for the date of extinction of the Thylacine in Tasmania. As per Table 1b (main paper), except based on all sightings rather than physical specimens and expert sightings only.

| | | McInerny et al. 2006 | | | Roberts & Solow 2003 | | |
|---|---|---|---|---|---|---|---|
| Scenario | n | MTE | 95% CI | PE-2020 | MTE | 95% CI | PE-2020 |
| Bioregional (all record types) | | | | | | | |
| South-East | 135 | 1996 | 1969–2021 | 0.952 | 2016 | 1976–2074 | 0.643 |
| North-East | 245 | 1998 | 1974–2022 | 0.951 | 2005 | 1975–2039 | 0.886 |
| North-West | 331 | 1998 | 1980–2017 | >0.999 | 2007 | 1985–2034 | 0.903 |
| West / World Heritage Area | 516 | 2008 | 1989–2020 | 0.935 | 2016 | 1993–2032 | 0.602 |

**Table S6**. Extinction date inference from the OLE model for individual bioregions, based on the combination of physical specimens and expert-sighting records.

| IBRA | n | MTE | 95% CI |
|---|---|---|---|
| Furneaux | 11 | 1979 | 1962–2023 |
| Central Highlands | 61 | 1985 | 1963–2008 |
| King | 109 | 1992 | 1967–2006 |
| Ben Lomond | 57 | 1996 | 1972–2026 |
| Southern Ranges | 82 | 1998 | 1956–2023 |
| Northern Slopes | 58 | 2005 | 1964–2033 |
| West | 121 | 2012 | 1970–2045 |
| South East | 16 | 2035 | 1921–2280 |
| Northern Midlands | 8 | 2069 | 1929–2310 |
| IBRA median | 58 | 1998 | 1963–2026 |
| Tasmania-wide | 528 | 2007 | 1988–2024 |

**Table S7**. Extinction date inference from the OLE model for individual bioregions, based on all available records.

| IBRA | n | MTE | 95% CI |
|---|---|---|---|
| King | 194 | 2002 | 1975–2032 |
| Central Highlands | 141 | 2003 | 1971–2047 |
| Ben Lomond | 191 | 2005 | 1975–2044 |
| Southern Ranges | 139 | 2008 | 1967–2052 |
| Northern Slopes | 137 | 2008 | 1977–2042 |
| West | 236 | 2016 | 1985–2040 |
| South East | 84 | 2023 | 1969–2197 |
| Furneaux | 54 | 2028 | 1965–2239 |
| Northern Midlands | 51 | 2104 | 1946–2716 |
| IBRA median | 139 | 2008 | 1971–2047 |
| Tasmania-wide | 1237 | 2016 | 1999–2026 |

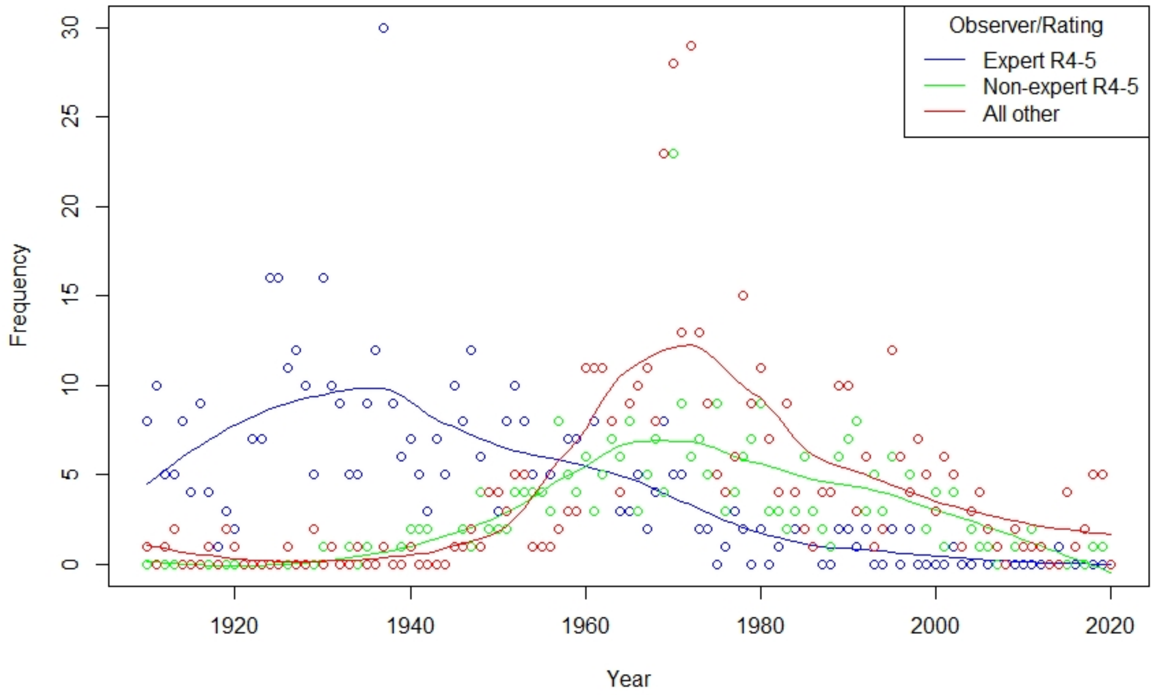


**Fig. S1**. Time series of Thylacine sightings in Tasmania, tallied by year, from 1910 through to 2019, and categorised as high quality (rating 4-5) for experts (blue) and non-experts (green), and all other sightings, rated 1-3 (red). The lines, separated by colour, are *loess* model fits (local polynomial regression smoother, with span = 0.75 and degree = 2).

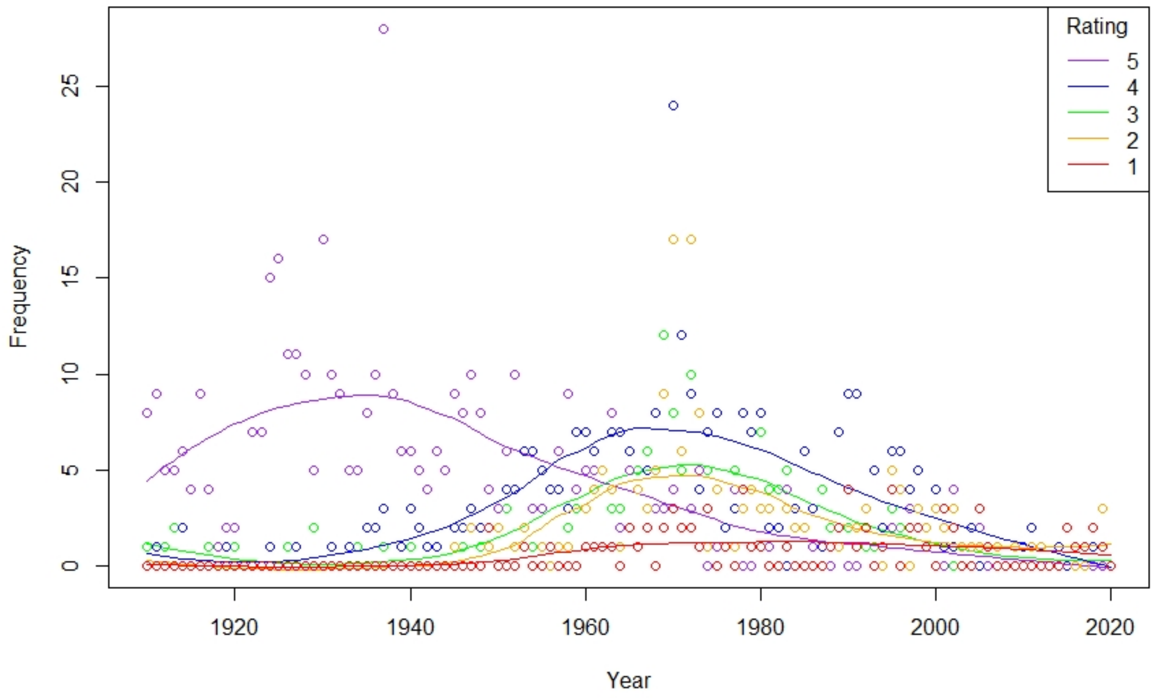


**Fig. S2**. Time series of Thylacine sightings in Tasmania, tallied by year, from 1910 through to 2019, and categorised as by quality rating, from 5 (highest) to 1 (lowest), as per Table S1. The lines, separated by colour, are *loess* model fits (see Fig S1).
